# High and low exogenous nitrate concentrations produce distinct calcium signatures in Arabidopsis roots

**DOI:** 10.1101/2025.03.03.641058

**Authors:** Stuti Shrivastava, Dilkaran Singh, Raymond E Zielinski, Amy Marshall-Colón

**Affiliations:** Department of Botany, University of Wisconsin Madison, Madison, WI, 53706, USA; Department of Plant Biology, University of Illinois Urbana-Champaign, Urbana, IL 61801, USA

## Abstract

Calcium (Ca^2+^) acts as a secondary messenger in plant responses to many stimuli, including nitrate (NO_3_ ^-^) signaling in roots. Nitrate uptake in plants occurs through two distinct transporter systems that have different affinities for the anion, which result in divergent downstream regulatory responses. However, it is unknown if the concentration dependent NO_3_ ^-^ response is linked to differential Ca^2+^ response. In this study, we used an Arabidopsis transgenic line expressing CBL1-mRuby2-GCaMP6s to investigate if there are unique Ca^2+^ signatures in intact root tissue in response to high and low NO_3_ ^-^ concentrations. A significant Ca^2+^ response was observed in root hairs in response to both high (5 mM) and low (0.25 mM) NO_3_ ^-^ concentrations. A distinct Ca^2+^ wave was observed in the low NO_3_ ^-^ treated roots, and an asynchronous Ca^2+^ response was captured in individual root epidermal cells. Cell-specific analysis revealed a distinct Ca^2+^ profile, comprising waves and spikes in response to NO_3_ ^-^, and prominent in root hairs but not in non-root hair bearing epidermal cells. The findings from this study demonstrate that there are distinct cell-specific and NO_3_ ^-^ condition-specific Ca^2+^ signatures in Arabidopsis roots.

## Introduction

Calcium (Ca^2+^) acts as a second messenger in response to multiple abiotic and biotic stimuli, such as cold, drought, light, pathogen, and fungal infections in all eukaryotic organisms (Riveras et al., 2015). The resting cytosolic concentrations of Ca^2+^ can range from 100 nM to 200 nM, while the Ca^2+^ reserves in the cell (e.g., vacuoles and free Ca^2+^ in the apoplast) can be several orders of magnitude higher, up to millimolar levels (Bender and Snedden, 2013; Medvedev, 2018; Wilkins et al., 2016). This large difference in the stored and resting concentration in the cell is maintained by active transport of Ca^2+^ from the cytosol into cellular organelles and the apoplast. Inwardly directed channels allow for a rapid and transient response to the stimuli generating Ca^2+^ flux within the organism. Briefly, the cytosolic free Ca^2+^ ([Ca^2+^]_cyt_) levels increase in response to the stimulus as the Ca^2+^ from subcellular compartments is released to the cytosol. The precise and unique response of stimulus-induced [Ca^2+^]_cyt_ is determined by the strength, duration, and frequency of the change in cytoplasmic concentrations and is referred to as the unique spatiotemporal ‘signature’ (Dodd et al., 2010; Wilkins et al., 2016). Changes in the [Ca^2+^]_cyt_ are captured by the Ca^2+^ signal sensor proteins, such as the well-characterized family of Ca^2+^ modulating protein or calmodulin (CaM), which play an important role in decoding the signature and relaying this information to other cellular proteins. The stimulus-specific Ca^2+^ signal is thus the result of this interplay between the Ca^2+^ signatures and the consequent Ca^2+^ sensing (Dodd et al., 2010). CaM-Ca^2+^ interactions with their target proteins were shown to have non-linear amplification of the Ca^2+^ signal through mathematical modeling of CaM binding transcriptional activators (CAMTA) (Virdi et al., 2015). Non-linear signal amplification is the reason behind the unique spatiotemporal signature of Ca^2+^ to different stimuli, and this fine-tuned response allows for much larger, but distinct downstream transcriptional responses in the cell.

Rapid Ca^2+^ signaling in response to nitrate (NO_3_^-^) uptake has been demonstrated in *Arabidopsis thaliana*. Arabidopsis plants expressing cytoplasmic aequorin (WT-AQ) protein and exposed to 5 mM NO_3_^-^ showed a transient increase in [Ca^2+^]_cyt_ within the first 10 sec in excised roots (Riveras et al., 2015). (2017) expressed GCaMP6 in Arabidopsis mesophyll cell protoplasts and observed a Ca^2+^ signal between 50 to 90 seconds in response to 10 mM NO_3_^-^. In addition, a Ca^2+^ signal in intact transgenic GCaMP6 seedlings exposed to 10mM NO_3_^-^ was shown in the root tip and the elongation zone of the root (Liu et al., 2017). The NO_3_ ^-^ concentrations used in these studies fall in the low affinity NO_3_ ^-^ uptake system (LATS) range (Liu et al., 2017; Riveras et al., 2015), characterized by NO_3_ ^-^ concentrations > 0.5 mM (Liu and Tsay, 2003). However, like many nutrient uptake systems, there is also a high affinity NO_3_ ^-^ uptake system (HATS) that is active at low NO_3_ ^-^ concentrations (< 0.2 mM for NO_3_ ^-^). There is evidence that a unique set of downstream target genes are modulated in response to either low or high [NO_3_ ^-^] in the soil (Bellegarde et al., 2017; Vidal et al., 2015). Thus, a reasonable hypothesis is that the different NO_3_ ^-^ uptake systems (HATS and LATS) elicit unique Ca^2+^ signatures. To date, there have been no quantitative studies investigating the Ca^2+^ signaling response to NO_3_ ^-^ concentrations in the high affinity range in intact plants. To address this knowledge gap, we treated whole Arabidopsis seedlings expressing a modified GCaMP6s Ca^2+^ biosensor with low (0.25 mM) or high (5 mM) NO_3_ ^-^ concentrations to capture high-resolution images of the resulting Ca^2+^ signatures using laser-scanning confocal microscopy. Our results support the notion that unique Ca^2+^ signatures are generated by low and high NO_3_ ^-^ that are most prominently observed in root hairs.

## Results

### Design of a novel Ca^2+^ sensor derived from GCaMP6s

In this study, we used an ultrasensitive GCaMP6s transgenic stable line in wild type (Col-0) Arabidopsis plants to study the real time Ca^2+^ signaling response in roots under different NO_3_^-^ conditions. The sensor was localized to the plasma membrane by incorporating 12 amino acids from the N-terminus of CBL1 (Batistic et al., 2008). In addition, ratio imaging of the signal generated by GCaMP6s binding of Ca^2+^ was facilitated by fusing the sensor to mRuby2. Figure S1 demonstrates that mRuby2 fluorescence in a purified, recombinant version of the sensor lacking the plasma membrane targeting sequence is minimally changed by resonance energy transfer from GCaMP6s and Ca^2+^. Further, since most cytosol in mature plant cells is closely appressed to the plasma membrane, a considerable proportion of any signal arising from increased [Ca^2+^]_cyt_ would be generated there.

### Low and high NO_3_^-^ trigger unique Ca^2+^ responses in Arabidopsis root hairs

Root hairs play an important role in nutrient acquisition and water uptake, as they provide a larger effective surface area that is in contact with the soil (Canales et al., 2017). To understand the signaling responses to NO_3_ ^-^ in root hairs (RHs), we treated the modified GCaMP6s Ca^2+^ biosensor Arabidopsis transgenic seedlings with low (0.25 mM) and high (5 mM) NO_3_ ^-^ solutions and captured the peak Ca^2+^ signal response in both RHs and non-root hair (NRH) bearing epidermal cells (Figure 1). Microscopy was focused on the maturation zone of the root where epidermal cells are fully differentiated and where RHs form, and where the sensitivity of the Ca^2+^ channel is greater than that of the cortex (Demidchik et al., 2007). When the seedlings were treated with low NO_3_ ^-^, a 3-fold increase (NO_3_ ^-^ to KCl) in Ca^2+^ signal was observed at peak intensity for RHs compared to a 2-fold increase (NO_3_ ^-^ to KCl) in the peak Ca^2+^ signal response for NRH cells (*p* = 0.03) (Figure 1A). Similarly, high NO_3_ ^-^ treatment resulted in a 2-fold increase (NO_3_ ^-^ to KCl) in the Ca^2+^ signal at peak intensity in RHs compared to a 1.25-fold reduced (NO_3_ ^-^ to KCl) Ca^2+^ signal observed in NRH cells (*p* = 0.001) (Figure 1B).

**Figure 1.**
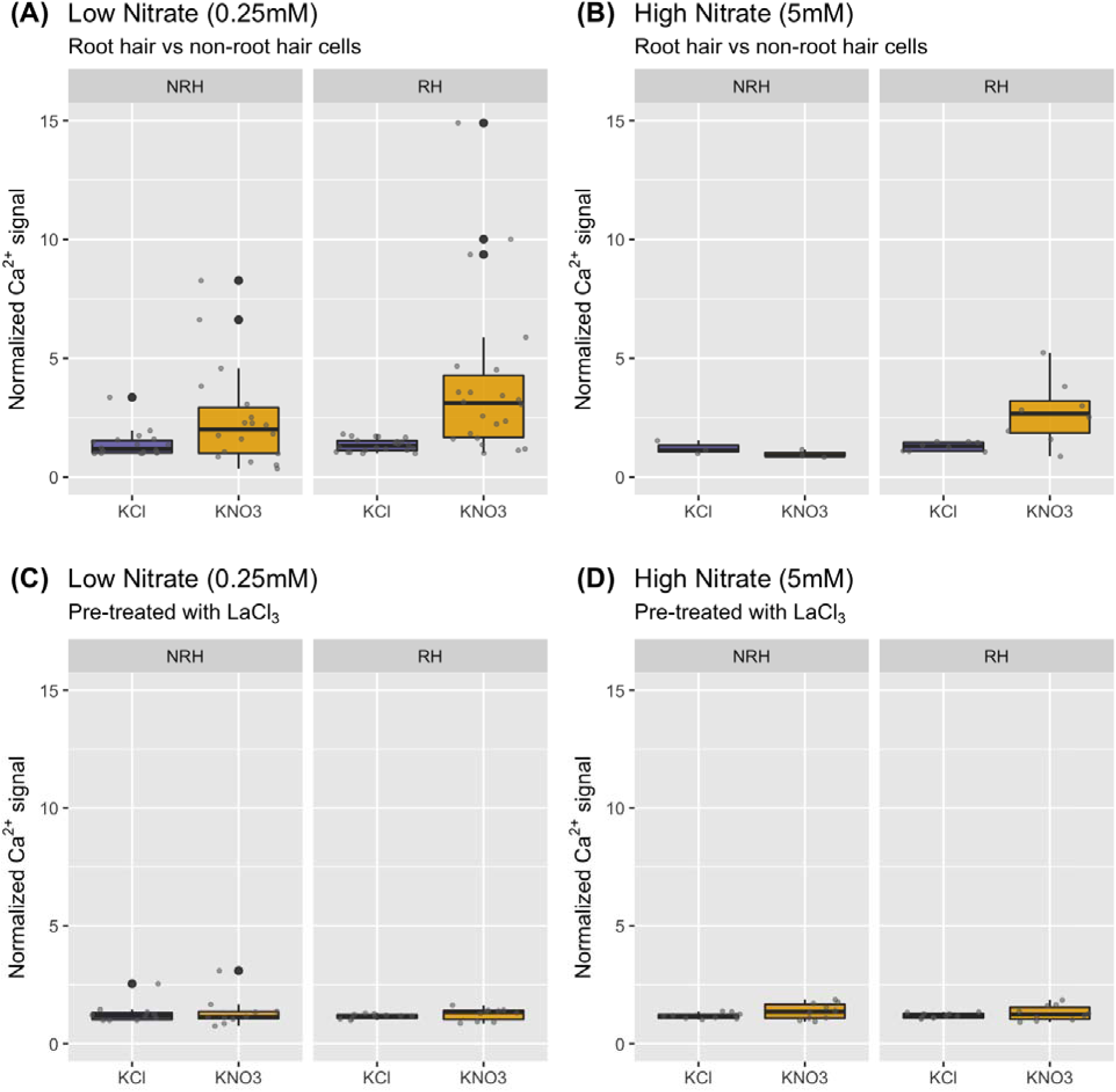
NO_3_^-^-induced increase in normalized Ca^2+^ signal at both low (0.25 mM) and high (5 mM) levels for root hair (RH; right panel in each pair) compared to non-root hair epidermal cells (NRH; left panel in each pair). The peak normalized Ca^2+^ signal intensity variation is depicted by the boxplot. **(A)** Variation in normalized Ca^2+^ signal in response to low (0.25 mM, n= 10 seedlings) NO_3_^-^ treatment **(B)** Variation in normalized Ca^2+^ signal in response to high (5 mM, n=2 seedlings) NO_3_^-^. **(C)** Variation in normalized Ca^2+^ signal in LaCl_3_ pretreated seedlings in response to low (0.25 mM, n= 5 seedlings) NO_3_^-^ treatment **(D)** Variation in normalized Ca^2+^ signal in in LaCl_3_ pretreated seedlings in response to high (5 mM, n=5 seedlings) NO_3_^-^ treatment. The median is represented by a solid line in the boxplot. Individual data points are shown as small grey points and outliers are large black points.

We investigated whether the Ca^2+^ signals observed were in response to NO_3_ ^-^ by pretreating the seedlings with 5 mM lanthanum chloride (LaCl_3_; a putative plasma membrane non-selective Ca^2+^ channel blocker) before replacing the solution with NO_3_ ^-^ treatment to prevent an increase in [Ca^2+^]_cyt_ levels in the plant cells (Grant et al., 2000). We performed all pretreatments for 1 hour to minimize the secondary effects on the cellular processes known to be associated with LaCl_3_ treatment (Tracy et al., 2008). The LaCl_3_ pretreatment abolished the unique Ca^2+^ response in the seedlings in both RHs and NRH cells, confirming the low NO_3_ ^-^-induced Ca^2+^ response (*p* = 0.53, Figure 1C) and high NO_3_ ^-^-induced Ca^2+^ response (*p* = 0.44, Figure 1D). These results demonstrate that a NO_3_^-^-induced Ca^2+^ signal occurs in response to both low- and high-NO_3_^-^ treatments in the RHs compared to NRH epidermal cells, and to a greater magnitude in response to low NO_3_^-^.

Both the RHs and NRH cells showed a dynamic Ca^2+^ profile over time for low and high NO_3_^-^ treatments (Figure S2A and S2B, Videos 1 and 2). We investigated whether the observed peak Ca^2+^ signal is a function of time by comparing the median time-to-peak Ca^2+^ signal intensity in both RHs and NRH cells. There was no significant association between the time-to-peak Ca^2+^ signal in the low NO_3_^-^ treatment in RHs despite a faster median time-to-peak Ca^2+^ response at 135 seconds compared to 160 seconds in the KCl treated seedlings. The median time-to-peak Ca^2+^ signal in NRH cells, irrespective of the treatment, was similar to RHs treated with KCl (Figure S2A). On the other hand, the median time-to-peak Ca^2+^ signal in response to high NO_3_^-^ treatment in the RHs was approximately 145 seconds after treatment compared to 350 seconds in KCl treated RHs (*p* = 0.045; Figure S2B). The median time-to-peak Ca^2+^ signal in the NRH cells showed an inverse trend compared to the RHs and was statistically insignificant. The KCl treated NRH cells responded 110 seconds after treatment, but the high NO_3_^-^ treated NRH cells had a median time-to-peak Ca^2+^ signal about 300 seconds after treatment (Figure S2B).

The variation in the time-to-peak Ca^2+^ signal and the magnitude of the peak Ca^2+^ response observed in RHs and NRH cells under both low- and high-NO_3_^-^ treatments demonstrate the asynchronicity of the short-term Ca^2+^ signal bursts that propagated through the root in response to a NO_3_^-^ stimulus. In this study, this asynchronous Ca^2+^ signal was most prominently observed in the RHs compared to the NRH bearing epidermal cells.

### Asynchronous Ca^2+^ signatures were observed in Arabidopsis root hairs in response to both low and high NO_3_^-^

The increase in [Ca^2+^]_cyt_ levels required for the generation of Ca^2+^ signals results in cellular concentrations that can be toxic (Medvedev, 2018). To prevent this lethal situation, a balance of influx and efflux carriers and channels regulates Ca^2+^ levels, appearing in short-term bursts, resulting in spikes, oscillations, and waves (Medvedev, 2018). Such short bursts were captured in this study by analyzing the Ca^2+^ signal response in adjacent sections of the RHs.

The RHs had a unique signal profile observed as a combination of Ca^2+^ spikes and waves under both low and high NO_3_^-^ treatments (Figure 2A and 2B, Videos 1 and 2). Under low NO_3_^-^, the root hairs had a 3.5- to 8-fold higher normalized Ca^2+^ signal (Figure 2A, panel RH-mid and RH-tip) compared to the NRH cell (Figure 2A, panel NRH). Moreover, the first Ca^2+^ spike was observed for the root hair sections at 190 seconds (Figure 2A, panel RH-tip, RH-mid, RH-base, Video 1) while a Ca^2+^ wave was observed for both the RH-base and RH-tip with peak amplitudes observed around 190 seconds and 250 seconds after low NO_3_^-^ treatment (Figure 2A, panel RH-base, Video 1).

**Figure 2.**
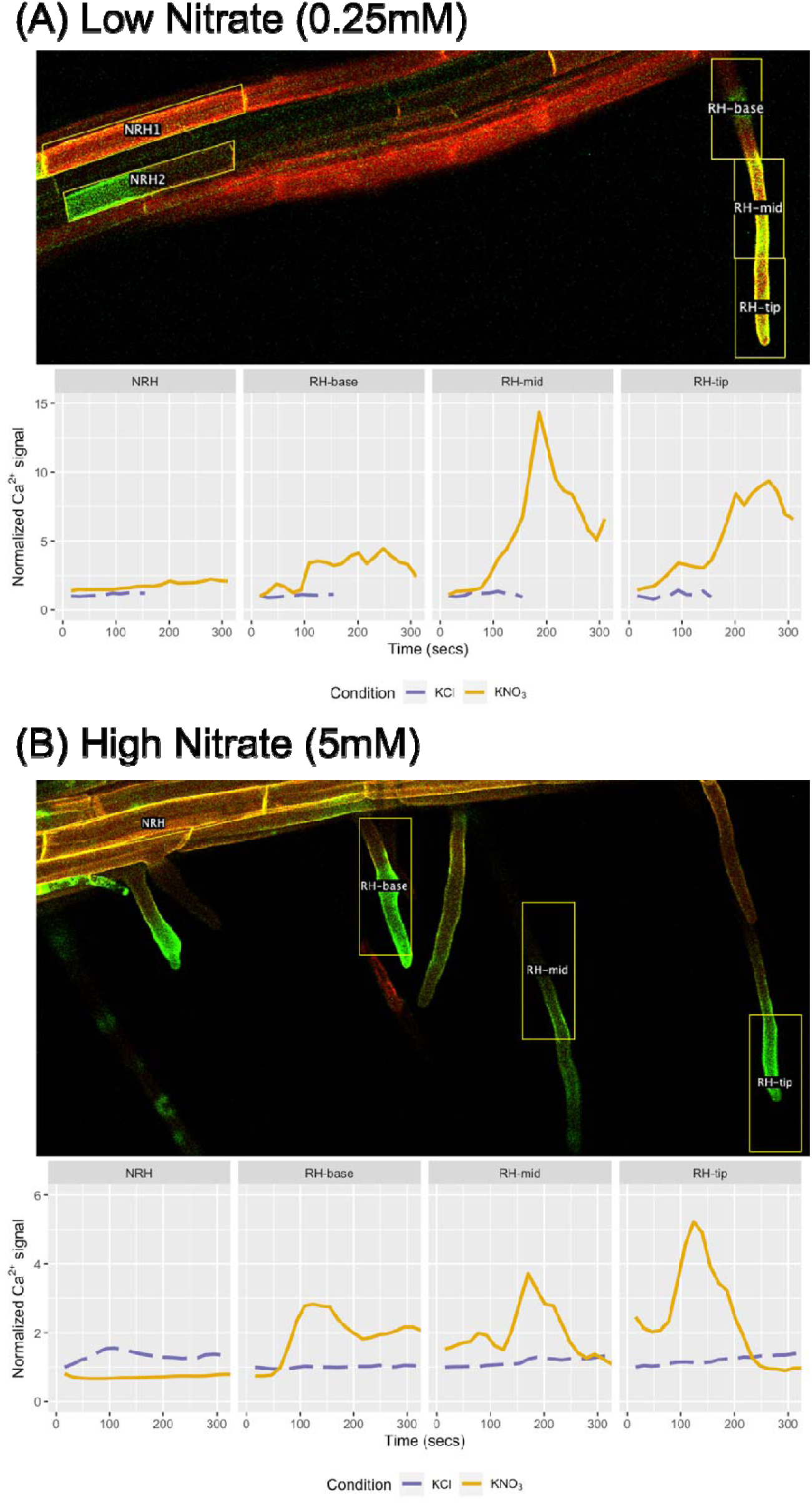
Asynchronous normalized Ca^2+^ signal responses of the root hair (RH) compared to non-root hair (NRH) bearing epidermal cells showing distinct signatures comprising waves and spikes under low (0.25 mM) and high (5 mM) NO_3_^-^ treatment. **(A)** The kinetics of apparent Ca^2+^ signal represented through the GFP fluorescence intensity responding to low NO_3_^-^ treatment. The root tissue imaged shows the apparent Ca^2+^ signal through GFP fluorescence with mRuby2 RFP background and captures the response 90 seconds after the NO_3_^-^ treatment. The peak normalized Ca^2+^ signal intensity was captured around 190 seconds for all RH sections (RH-base, RH-mid and RH-tip) while NRH cell had peak normalized Ca^2+^ signal intensity was around 240 seconds after low NO_3_^-^ treatment. **(B)** The kinetics of apparent Ca^2+^ signal responding to high NO_3_^-^ treatment for the selected root sections. The root tissue imaged shows the apparent Ca^2+^ signal through GFP fluorescence with mRuby2 RFP background and captures the response 110 seconds after the NO_3_^-^ treatment. The peak normalized Ca^2+^ signal intensity was captured around 120 seconds for RH-tip, and close to 140 seconds for RH-mid and RH-base sections.

Under high NO_3_^-^, the emerging root hair had a Ca^2+^ wave with first peak wave amplitude from 110 to 155 seconds, followed by a second wave with peak amplitude between 280 and 325 seconds (Figure 2B, panel RH-base). In contrast, both the tip of the mature RH and the mid-section of the RH revealed a Ca^2+^ spike with peak amplitudes around 120 and 170 seconds after treatment, lasting 125 and 95 seconds respectively (75 to 200 seconds for RH-tip and 140 to 235 seconds for RH-mid section; Figure 2B, panels RH-tip and RH-mid, Video 2). In comparison, the Ca^2+^ signal was much lower and remained unchanged over the same time frame in NRH cells (Figure 2B, panel NRH). At its peak amplitude, the Ca^2+^ signal in the RH showed a 4- to 7.5- fold increase compared to the NRH cell treated with high NO_3_ ^-^ at the same time after treatment (Figure 2B, panel RH-base and RH-tip).

The dynamic Ca^2+^ signatures captured post NO_3_ ^-^ treatment reveal asynchronous, short-term Ca^2+^ signal bursts, which propagate through the RHs in response to the NO_3_ ^-^ stimulus. Root hair Ca^2+^ signatures appear to change as the RHs mature. Growing root hairs usually display frequent, shorter Ca^2+^ waves while the mature root hairs show Ca^2+^ spikes from mid-RH to tip of the RH with high signal intensity (Figure 2).

### Low but not high NO_3_ ^-^ transiently induces cytosolic Ca^2+^ response in bulk cell analysis of roots

Nitrate as a signaling molecule triggers both cell-specific and tissue-specific responses, thereby initiating a cascade of both local and systemic signaling in plants (Vega et al., 2019). To investigate the systemic response in epidermal cells, we performed a bulk analysis of the various segments captured across the length of the root maturation zone, whereas the above results sections describe single cell-type analyses. When the maturation zone of the root was analyzed for the tissue-specific NO_3_^-^ response, low NO_3_^-^ treatment revealed a 2-fold increase in the Ca^2+^ signal compared to the KCl treatment (*p* = 3.58e-11, Figure 3A). The absence of this unique Ca^2+^ response in the LaCl_3_ pretreated seedlings confirmed that low NO_3_^-^ treatment triggered the increase in [Ca^2+^]_cyt_ (*p* = 0.19, Figure 3C). We observed a significant Ca^2+^ signal response for the interaction of low NO_3_^-^ treatment and time, which can be explained by the washout of the inhibitor from the plant cells (*p* = 0.013). The median time-to-peak Ca^2+^ signal was not significant in LaCl_3_ pretreated seedlings (*p* = 0.21). Thus, similar to the cell-specific response to low NO_3_^-^ treatment in RH vs NRH cells, an increase in [Ca^2+^]_cyt_ occurs in the bulk cell analysis of the maturation zone as well. This suggests a homogenous Ca^2+^ response across epidermal cells in the maturation zone in response to low NO_3_^-^.

**Figure 3.**
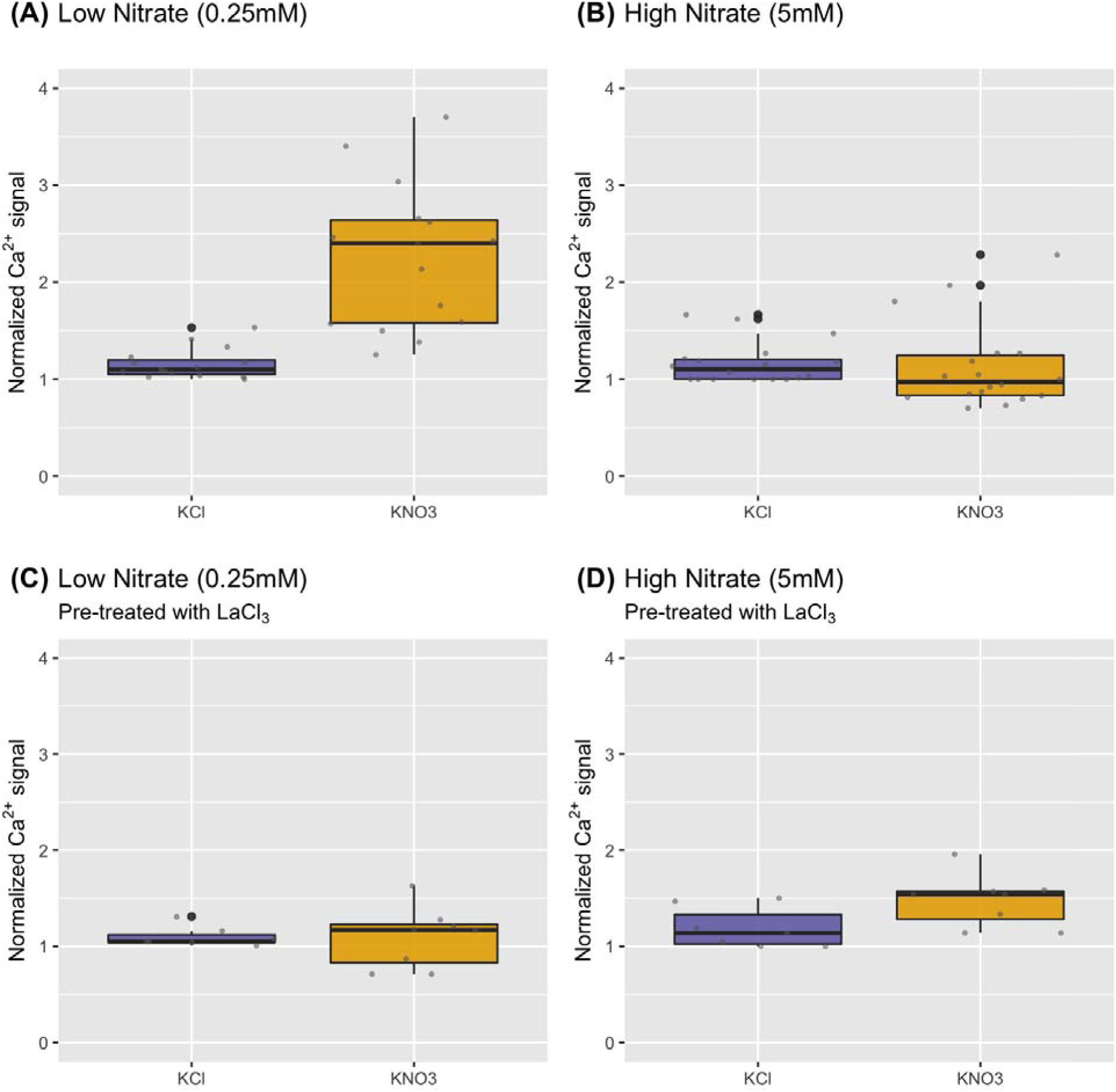
NO_3_^-^-induced peak normalized Ca^2+^ signal shows an increase in low (0.25 mM) NO_3_^-^ treatment but not in high (5 mM) treatment in epidermal cells in the root maturation zone. For pretreatment, the seedlings were incubated with 5mM LaCl_3_ for 1 hour. **(A, B)** Variation in the peak normalized Ca^2+^ signal in response to low (0.25 mM, n= 5 seedlings) NO_3_^-^ and high (5 mM, n=7 seedlings) NO_3_^-^ treatment. **(C, D)** Variation in the peak normalized Ca^2+^ signal in response to low (0.25 mM, n= 6 seedlings) NO_3_^-^ and high (5 mM, n=7 seedlings) NO_3_^-^ treatment LaCl_3_ pretreated seedlings. The median is represented by a solid line in the boxplot. Individual data points are shown as small grey points and outliers are large black points.

In contrast to the cell-type specific response described in the above sections, no unique Ca^2+^ signatures were observed between treatment with high NO_3_ ^-^ and the KCl control (*p* = 0.11, Figure 3B) in the bulk cell analysis across the root maturation zone. To explore the possibility of other stressors eliciting the Ca^2+^ signal, we pretreated the seedlings with LaCl_3_ for 1 hour before replacing the pretreatment solution with 5mM KNO_3_ or KCl. No significant difference was observed between KCl control and high NO_3_ ^-^ treatment in the LaCl_3_ pretreated seedlings (*p* = 0.59, Figure 3D), and, unlike low NO_3_ ^-^ treatment, there was no significant NO_3_ ^-^ treatment by time interaction (*p* = 0.68, Figure 3D) at high [NO_3_ ^-^]. Further, there were no statistically significant difference observed for the median time–to-peak Ca^2+^ signal under high NO_3_ ^-^ treatment (Figure S3). The discrepancy between the Ca^2+^ response to 5 mM NO_3_ ^-^ in single vs bulk epidermal cell analysis suggests that the Ca^2+^ response to high NO_3_ ^-^ is heterogeneous, where it is more prominently observed in RHs compared to the NRH epidermal cells.

### Dynamic Ca^2+^ signatures observed in the roots in response to low NO_3_ ^-^

The Ca^2+^ signatures observed as short bursts in the RHs were also captured in this study at the tissue level in the bulk cell analysis of subsequent sections of the root maturation zone in response to low NO_3_ ^-^ treatment. We observed an asynchronous Ca^2+^ wave in the root tissue after low NO_3_ ^-^ treatment, with a 2.5- to 3.5-fold increase of the Ca^2+^ signal compared to the KCl control (Figure 4, panels S1 and S2). However, under high NO_3_ ^-^ treatment, no distinct Ca^2+^ signature was observed at the tissue level (Figure S4).

**Figure 4.**
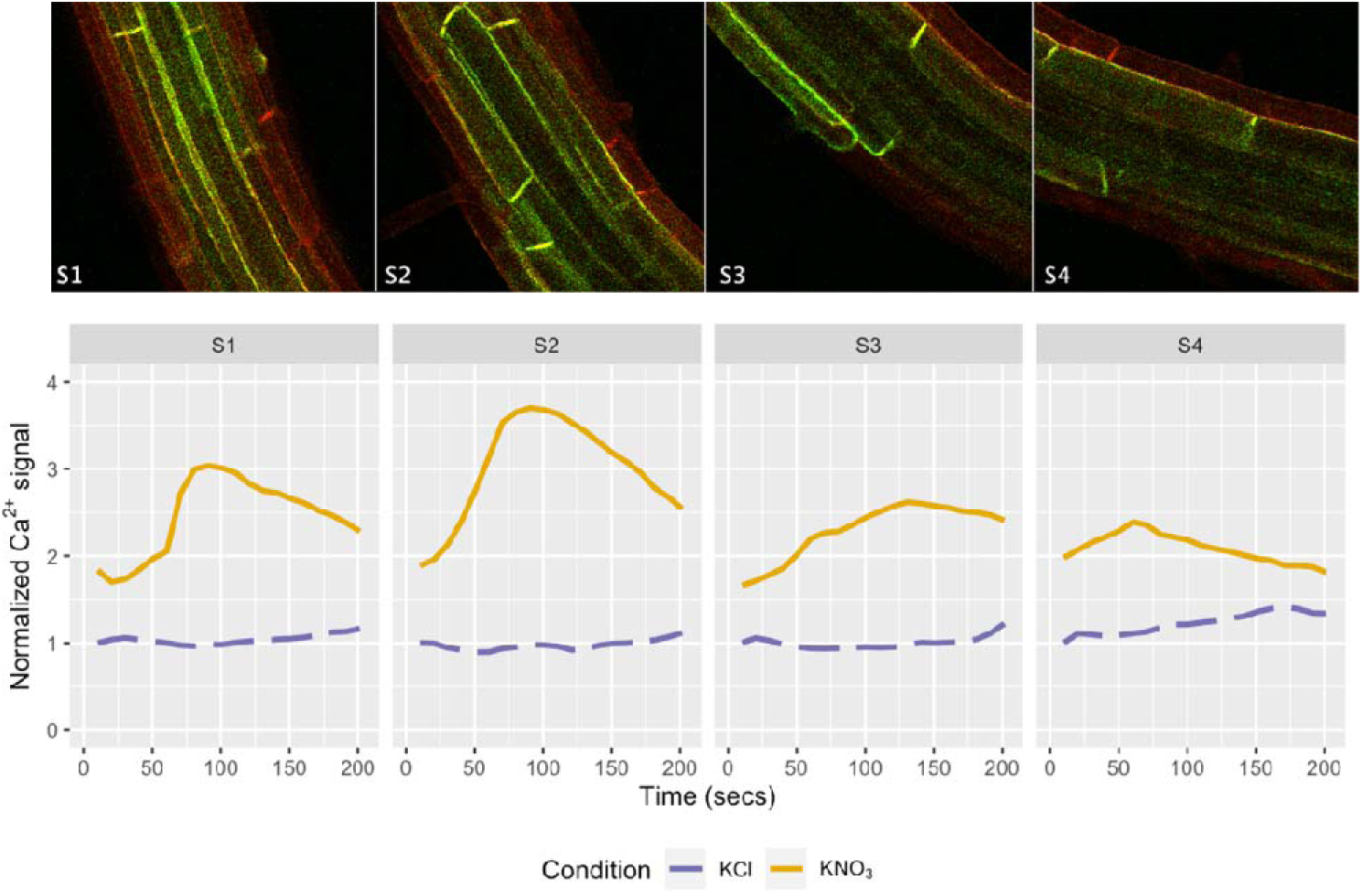
Asynchronous normalized Ca^2+^ signal in the root sections in response to the low NO_3_^-^ treatment. A representative root section for the low NO_3_^-^ treated seedling is shown with the normalized Ca^2+^ signal, where section S1 was distal to the root tip while S4 was proximal to the root tip. The root tissue imaged above shows the apparent Ca^2+^ signal through GFP fluorescence with mRuby2 RFP background and captures the response 120 seconds after low NO_3_^-^ treatment. The peak normalized Ca^2+^ signal intensity was captured around 90 seconds for sections S1 and S2, while the peak intensity of the normalized Ca^2+^ signal was close to 125 seconds for S3 and 60 seconds for S4.

## Discussion

Roots provide structural support and access to the nutrients and water necessary for plant survival and growth. Roots and root hairs (RHs) perceive the exogenous nitrogen nutrient status as signals. Several NO_3_^-^ transporters are localized in roots and root hairs (RHs), such as NPF6.3, NRT2.1, and NRT2.6 (Canales et al., 2017), among others. Both RHs and NRH cells can perceive signals of NO_3_^-^ status and trigger a signaling response that dynamically regulates root system architecture or defense responses (Wang et al., 2018). In tomato, spinach, oilseed rape, and four grass species, both RH density and RH length were negatively correlated with increasing NO_3_^-^ concentration (B. Liu et al., 2020). On the other hand, RH density increased in several ecotypes of Arabidopsis in response to an increase in local NO_3_ ^-^ levels (Vatter et al., 2015) suggesting that both local and systemic NO_3_^-^ signals play a role in RH development. Nitrate-induced Ca^2+^ signals trigger the primary NO_3_^-^ response, resulting in the transcriptional regulation of NO_3_^-^ uptake and assimilation, and lateral root and RH development (Canales et al., 2017; B. Liu et al., 2020). While the high- and low-affinity NO_3_^-^ transporter systems and their related down-stream signaling are well-studied in plants (Fredes et al., 2019), the relationship between these two systems and Ca^2+^ signaling is unknown. In this study, we addressed this knowledge gap by examining Ca^2+^ signal responses in differentiated epidermal root cells to low and high NO_3_^-^ concentrations in the known ranges for the HATS and LATS systems, respectively. We achieved this by using an ultrasensitive modified GCaMP6s transgenic line in Arabidopsis to investigate the Ca^2+^ signaling dynamics in roots in response to NO_3_^-^. The Hybriwell chamber allowed us to capture high-resolution Ca^2+^ signal responses observed in the roots of the seedlings placed in treatment solution, specifically in the NRH bearing epidermal cells and the RHs themselves. But placing the seedling in solution and not fixed on a slide also made it difficult to capture the Ca^2+^ responses of the RH bearing epidermal cells and the RHs themselves in the same focal plane. For this purpose, we compared the Ca^2+^ signals between the RHs and the NRH bearing epidermal cells in this study.

Both high and low NO_3_^-^ treatments elicited an increase of 2- to 3-fold (NO_3_^-^ to KCl), respectively, in the Ca^2+^ signal in the RHs compared to NRH cells (Figure 1). Pretreatment with LaCl_3_, a non-selective putative plasma membrane Ca^2+^ channel blocker, revealed that the Ca^2+^ response was specific to NO_3_^-^ treatment. The observed Ca^2+^ response to high [NO_3_^-^] in RHs agrees with those of previous studies in Arabidopsis mesophyll protoplasts (Liu et al., 2017) and excised roots (Riveras et al., 2015), but we report the first measurement of the Ca^2+^ response to low [NO_3_^-^]. At coarse resolution (bulk cell analysis), a Ca^2+^ response was only observed in response to low NO_3_^-^ treatment (Figure 3). This demonstrates that there is a stronger and potentially a more homogenous Ca^2+^ response to treatment with low [NO_3_^-^] compared to high [NO_3_^-^]. Our results show a dynamic and unique Ca^2+^ response to variable NO_3_^-^ concentrations in intact root tissue, which are most clearly observed in RHs in response to high NO_3_^-^. The absence of a distinct Ca^2+^ response to high NO_3_ ^-^ treatment in the bulk cell analysis across the maturation zone of the root could be due to a dampening effect caused by differing kinetics of individual cell types in the root. Liu et al. (2017) quantified a significant Ca^2+^ response to 10 mM NO_3_^-^ in a uniform population of shoot mesophyll protoplasts expressing GCaMP6, partially supporting our findings. Protoplasts lack connections to other neighboring cells so can be used to report individual cell type responses, but are inherently unable to capture intercellular responses, such as the dynamic process of cell-to-cell Ca^2+^ signaling (Pasternak et al., 2020).

To further support the notion of differing Ca^2+^ kinetics in individual cell types, we observed an asynchronous response to NO_3_^-^ stimulus in the root epidermal cells, quantified as the variation in the median time-to-peak Ca^2+^ signals (Figure S2). A faster peak Ca^2+^ response was observed in the low NO_3_^-^ treated seedlings compared to the KCl treatment, but high NO_3_^-^ treatment induced a statistically significant median time-to-peak response that was 200 seconds faster than median time-to-peak response to the corresponding KCl treatment. The median time to a Ca^2+^ spike in the RH tip was observed close to 120 seconds in the high NO_3_^-^ treatment but was much slower in low NO_3_^-^ treatment at approximately 190 seconds after treatment. The response time of NO_3_^-^-induced Ca^2+^ signal to high NO_3_^-^ in both RH and root tissue is similar to that observed previously in mesophyll cells, despite the differences in the two systems (K.-H. Liu et al., 2020; Liu et al., 2017). Our analysis provides the first quantitative evidence in intact root tissue in GCaMP6s expressing Arabidopsis plants that account for the asynchronicity observed in response to the NO_3_^-^ treatments.

In the present study, unique Ca^2+^ waves were observed in the low NO_3_^-^ treatment in the bulk cell (Figure 4) and the cell type-specific comparison between RH and NRH cells (Figures 2 and 4). The peak amplitude of the Ca^2+^ signal varied between 3- to 8-fold increase in the RHs for both high- and low- NO_3_^-^ treatments, and a 3-fold increase in the whole root response to low NO_3_^-^ treatment. The unique Ca^2+^ signatures also showed that the growing root hairs had frequent shorter waves, while the mature root hairs had single spikes with high signal intensity towards the tip of the root hair (Figure 2). The Ca^2+^ signal observed for the RHs in response to both low- and high- NO_3_^-^ treatment in this study had a profile similar to the transient Ca^2+^ signals observed when auxin was introduced in individual root cells using a constant low current (iontophoretically) (Dindas et al., 2018). Interestingly, the auxin induced RH tip gradient is responsible for signaling RH growth and elongation (Wymer et al., 1997). Likewise, rapid Ca^2+^ waves have been observed in response to salt stress, which were long-distance signals traversing the length of an Arabidopsis seedling, relaying the information to the whole plant (Choi et al., 2014). Since rapid information transmission does occur in plants to relay information from the roots to shoots, it is reasonable to suggest that the NO_3_ ^-^ dependent Ca^2+^ signal is similarly long-distance encoded information. Further work is needed to explore the cell-type specific Ca^2+^ signals in RHs to understand the mechanism of NO_3_ ^-^-responsive RH growth to NO_3_ ^-^ acquisition. Additionally, investigating the root-to-shoot response would lead to a better understanding of the root-shoot-root relay observed in response to NO_3_ ^-^ at the level of gene expression (Bellegarde et al., 2017).

## Conclusions

In conclusion, this study utilized an ultrasensitive Ca^2+^ biosensor GCaMP6s to elucidate the patterns of Ca^2+^ response to variable NO_3_ ^-^ treatments. The NO_3_ ^-^ transporter protein, NPF6.3, functions as a nutrient transporter as well as a sensor receptor – a transceptor (Giehl and von Wirén, 2015; Vidal et al., 2020). The response to NO_3_ ^-^ as a signaling molecule was identified through Ca^2+^ ions acting as the second messenger in the NO_3_ ^-^ signaling pathway (Riveras et al., 2015). The presence of two different NO_3_ ^-^ uptake kinetics in plants, each responding to NO_3_ ^-^ and triggering a distinct set of downstream responses suggested that the NO_3_ ^-^ signal is being uniquely perceived at different concentrations and trigger distinct downstream responses (Bellegarde et al., 2017; Riveras et al., 2015). The low concentration NO_3_ ^-^ treatment (0.25 mM) triggers the HATS uptake activity, and at this NO_3_ ^-^ concentration, we observed a unique and transient rise in [Ca^2+^]_cyt_ in Arabidopsis roots. A Ca^2+^ wave was observed in seedlings treated with low [NO_3_ ^-^], which suggests a possible long-distance transmission of information, similar to observations made during salt stress that generated a long distance Ca^2+^ wave (Choi et al., 2014). A stronger Ca^2+^ response was captured in RHs compared to the NRH cells under both low and high NO ^-^ treatments, with short bursts of Ca^2+^ signals observed as waves and spikes. These short bursts triggered in response to NO ^-^ are similar to the transient signals observed in response to auxin (Dindas et al., 2018); however, further work is required to understand the role of Ca^2+^ response to NO_3_ ^-^ in RH elongation. Together, these results highlight distinct Ca^2+^ signatures that occur in response to low and high NO_3_ ^-^ in the RHs, that occur as Ca^2+^ waves and spikes, and suggest that the asynchronous Ca^2+^ signatures observed in root epidermal cells may relay information across the root tissue, at least in response to low NO_3_ ^-^.

## STAR Methods

### Plasmid construction

A binary plasmid harboring the sequence encoding CBL1-mRuby2-GCaMP6s was constructed as outlined in Figure S1. Sequences encoding mRuby2 (#40260, (Lam et al., 2012) and GCaMP6s (#40753, (Chen et al., 2013) were obtained from Addgene (Watertown, MA). Synthetic oligonucleotides encoding the N-terminal 12 amino acids of CBL1 (At4g17615, (Batistic et al., 2008) were purchased from Eurofins Genomics (Louisville, KY). The CaMV35S promoter and Tobacco etch virus (TEV) 5’ translational enhancer sequences were derived from pRTL2 (Carrington et al., 1991). The sequences were assembled in pGreenII 0179 (Hellens et al., 2000). The plasmid was introduced into *Agrobacterium tumefaciens* GV3101 (pMP90) and the resulting cells were used to transform *Arabidopsis thaliana* Col-0 by a modified floral dip protocol (Davis et al., 2009). Transgenic seedlings were selected on ½ strength Murashige and Skoog Basal media containing 1% (w/v) sucrose, 2.5mM MES, pH to 5.7 with KOH, and 20 μg/ml hygromycin. Homozygous individuals were selected following three generations of selection for hygromycin resistance.

### Plant material

*Arabidopsis thaliana* ecotype Columbia-0 (Col-0) seedlings genetically modified to express the Ca^2+^ biosensor (p35S-CBL1-mRuby2-GCaMP6s) were used in this study. All seeds were surface sterilized with 75% ethanol solution and stratified in sterile autoclaved nuclease-free (ANF) water for a minimum of 48 hours at 4°C in dark before sowing to synchronize germination. Stratified seeds were sown on ½ strength Murashige and Skoog Modified Basal Salt mixture without Nitrogen (MS -N; M531, PhytoTechnology Laboratories, Lenexa, KS) supplemented with 2.5 mM ammonium succinate, 2.5 mM MES, 1% (w/v) sucrose, 2% (w/v) agar, and pH to 5.7 with NaOH. All seedlings were grown in vertical plates under long day (16 hours day/ 8 hours night) at 22°C and 120 μmol m^-2^ s^-1^ of photosynthetically active radiation (PAR) for 3 days before imaging.

### Mounting samples in Hybriwell™ and treatments

The Hybriwell™ sealing system HBW20 (Grace BioLabs, Oregon, USA) was used to securely seal the 3-day-old seedlings on to cover glass 50mm x 35mm (Fisher Scientific, Waltham, MA) with autoclaved, nuclease-free (ANF) water (pH ∼5.6) as described previously (Vang et al., 2018). Briefly, the seedling was first placed on the cover glass in about 10 µL of ANF water drop and the Hybriwell™ chamber was placed gently while minimizing air bubbles. An additional 200 µL of ANF water was pushed in through one port to remove all air bubbles, which forced water to eject out of the other port. Lastly, 100 µL ANF water drops were placed on both the ports to prevent evaporation of the 30 µL water in the chamber. The sealed seedlings on the cover glass were placed in sterile square plates to maintain humidity and sterility and were incubated in the growth chamber under conditions as described above, laying horizontally for at least 1 hour to minimize stress responses due to mounting of plant samples before the treatment and imaging. Two different NO_3_^-^ treatment conditions of 0.25 mM KNO_3_ (pH 5.0) and 5 mM KNO_3_ (pH 4.8) were used to study HATS and LATS responses respectively, with corresponding KCl concentrations as control (pH 4.9 and pH 4.8 for 0.25 mM and 5 mM KCl, respectively). Abiotic and biotic stressors can trigger an increase in cytoplasmic Ca^2+^ (Choi et al., 2014), which was blocked by using lanthanum chloride (LaCl_3_), a cytoplasmic Ca^2+^ channel blocker. Once the seedlings were sealed in Hybriwell™ on the cover glass, they were incubated for 1 hour in the growth chamber with 5 mM LaCl_3_ (pH 4.6) before further treatments.

### Real-time [Ca^2+^] imaging

Homozygous Arabidopsis transgenic lines expressing the Ca^2+^ sensor, *CBL1-mRuby2- GCaMP6s* under CaMV35S promoter, mounted and equilibrated as described above, were imaged using an EC Plan-Neofluar 10X/0.3 objective on a Zeiss LSM 710 multi-photon confocal microscope (Zeiss, Obercohen, Germany). The following settings were used on LSM710 to obtain images: pixel dwell: 1.024 µs, average: line 2; master gain: 700; pinhole diameter: 39.5 µm; filters: 489-647 nm; beam splitter: MBS 488/561; lasers: 488 nm (Argon 488) and 561 nm (DPSS 561-10). The mRuby2 sensor was excited with 561 nm and emission was captured between 560 nm and 647 nm, while the GFP sensor had 488 nm excitation and emission between 489 and 560 nm. The fluorescence intensities were acquired simultaneously by using a line scan average. The root images were first acquired as baseline pretreatment. The control solution was pipetted in through one of the ports on the Hybriwell™ unit while using a Kimwipe to wipe excess solution ejecting through the other port. The assumption was made that once the solution was pipetted in, any previous solution moved out and the resulting 30 µL of the solution in the chamber is homogenous. The time-series experiment was initiated where the images were captured every 10-25 sec for 3-10 mins. The same plant was then treated with the respective NO_3_ ^-^ solution and the time series experiment captured images at equally spaced time points as before. High resolution 1024×1024 pixel 12-bit images were captured for the root sections.

### Image analysis

The confocal images were analyzed using the FIJI analysis package (Schindelin et al., 2012) with the BioFormats plugin (Linkert et al., 2010), a distribution of ImageJ. The split channel setting was used to extract the fluorescent intensity levels of GFP and RFP from each confocal image. The mean fluorescence intensity was obtained for both RFP and GFP signals by using polygon region of interests (ROIs) across the root slices for each time point. For the cell-specific analysis, the polygon ROIs were manually assigned for multiple similar sized cells across the root image slice for each time point. A similar area of rectangular ROI was selected as background, which was subtracted from all other ROIs for each time point for each image.

The intensity data was exported for each channel and all further calculations were performed in Microsoft Excel. The Ca^2+^ signal was normalized in two steps. First, the proportionality of the Ca^2+^ signal (*F_i_*) was calculated as *F_i_* = *F_g_/F_r_*, where *F_g_* is the GFP (Ca^2+^) signal and *F_r_* is the invariant mRuby2 control signal. Then the Ca^2+^ signal was normalized with respect to the time 0 for KCl treatment, resulting in a normalized Ca^2+^ signal with a value of 1.0 for KCl time 0. This normalized Ca^2+^ signal is reported as ‘Ca^2+^ signal’ in this study for clarity. At least 5 biological replicates were used for all instances unless specified otherwise. The videos were generated using the cropped images in ImageJ.

### Statistical testing

A linear mixed effects analysis was performed for all datasets using the ‘lme4’ package in R (Bates et al., 2015; R Core Team, 2021). The fixed effects of NO_3_^-^ treatment (condition) and time were used with interaction for non-root hair specific datasets. The fixed effect of root hair (root hair vs non-root hair cell) was added with interaction for root hair specific datasets. The random effect was added as intercept for plant (subject) for all datasets. The contrasts were obtained using *contr.sum* function from ‘stats’ package in R. In all cases, the full model (described above) was compared with the reduced model (without the time as factor) using *anova* function from ‘stats’ package in R. The reduced model was used where no statistically significant information was added by the complete model to explain the data. Due to the unbalanced design of the sample datasets, all model significance outcome was obtained using the *Anova* function in ‘car’ package with Type III sum of squares (Fox and Weisberg, 2019). The assumptions for ANOVA analysis were checked using the Shapiro-Wilk Normality test and the Kolmogorov-Smirnov test from ‘stats’ package in R. The median test was used from the ‘agricolae’ package in R as the post hoc nonparametric test that applies chi-square test to test the probability of error that samples are independent (Felipe de Mendiburu, 2021).

## Author Contributions

Conceptualization and Methodology, S.S., A.M.C., R.E.Z; Investigation and Formal Analysis, S.S. and D.S.; Writing – Original Draft, S.S., A.M.C., R.E.Z.; Writing – Review & Editing, S.S., A.M.C., R.E.Z., and D.S.; Funding Acquisition and Supervision, A.M.C.

## Acknowledgments

The authors would like to thank Dr. Simon Gilroy and Dr. Nathan Miller at the University of Wisconsin, Madison for valuable discussions and advice on statistical analysis of the data.

## Video Legends

**Video 1.** Asynchronous NO_3_^-^-induced normalized Ca^2+^ response to low (0.25 mM) NO_3_^-^ treatment in the root hairs (RHs) and the non-root hair (NRH) bearing epidermal cells in the root maturation zone. A representative root section for low NO_3_^-^ treated seedling was selected to show the apparent Ca^2+^ signal represented through the GFP fluorescence intensity. The top panels show the KCl treatment response while the bottom panels show the NO_3_^-^ treatment. The left panel shows the RH response, and the right panel shows the NRH response in the representative sample.

**Video 2.** Asynchronous NO_3_^-^-induced normalized Ca^2+^ response to high (5 mM) NO_3_^-^ treatment in the root hairs (RHs) and the non-root hair (NRH) bearing epidermal cells in the root maturation zone. A representative root section for high NO_3_^-^ treated seedling was selected to show the apparent Ca^2+^ signal represented through the GFP fluorescence intensity. The top panels show the KCl treatment response while the bottom panels show the NO_3_^-^ treatment. First three panels from the left show the RH response, while the right panel shows the NRH response in the representative sample.

