## Supplemental Figures for "High and low exogenous nitrate concentrations produce distinct calcium signatures in Arabidopsis roots"

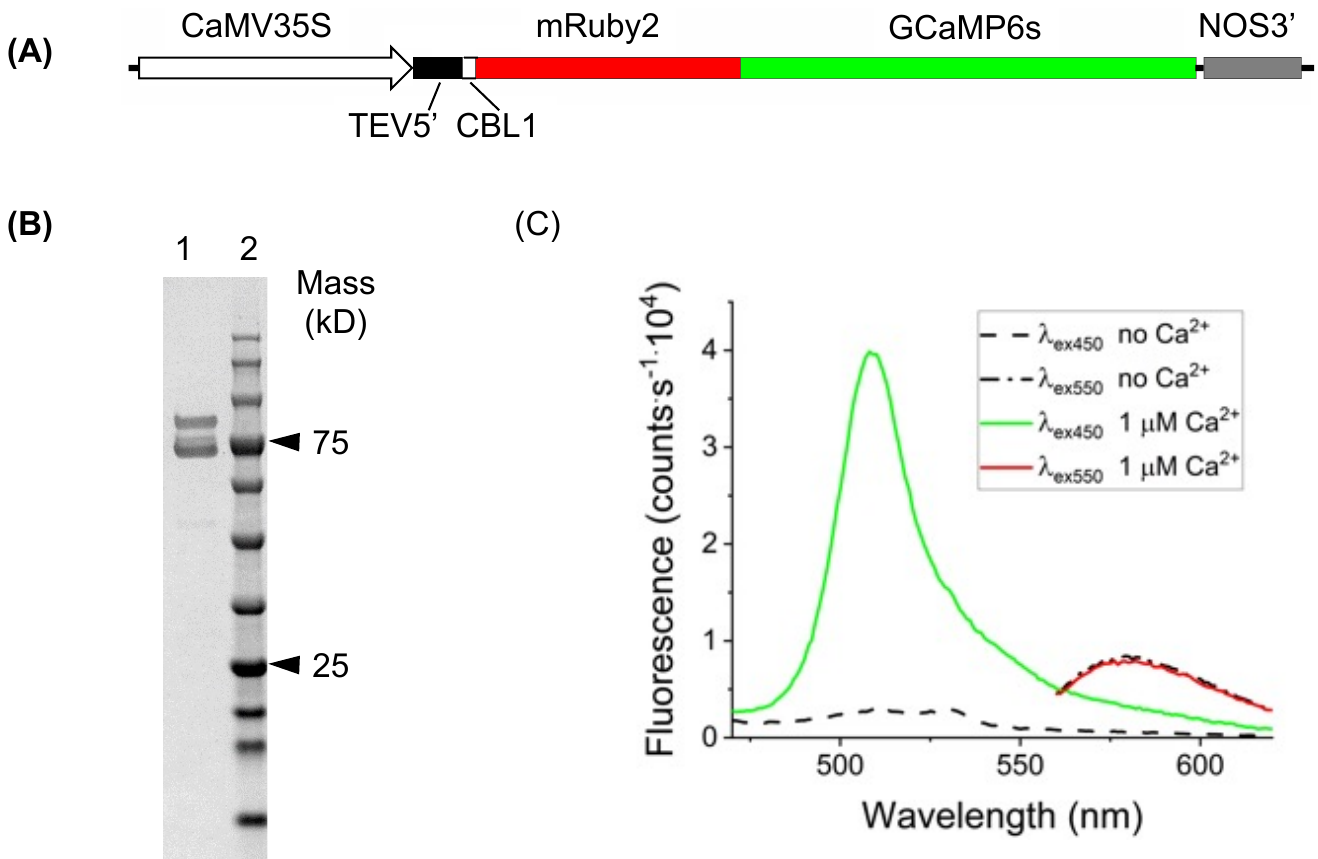

**Figure S1.** Design and functional characteristics of the modified GCaMP6s Ca^2+^ sensor. **(A)** Physical map of the construct introduced into *Arabidopsis thaliana* Col-0 plants. Sequences encoding mRuby2 (#40260, Lam et al., 2012) and GCaMP6s (#40753, Chen et al., 2013)) were obtained from Addgene (Watertown, MA). Synthetic oligonucleotides encoding the N-terminal 12 amino acids of CBL1 (At4g17615, Batistic et al., 2008) were purchased from Eurofins Genomics (Louisville, KY). The CaMV35S promoter and Tobacco etch virus (TEV) 5’ translational enhancer sequences were derived from pRTL2 (Carrington et al., 1991). The sequences were assembled in pGreenII 0179 (Hellens et al., 2000). **(B)** SDS-PAGE of recombinant 8X His-mRuby2-GCaMP6s. A His-tagged version of the modified GCaMP6s sensor lacking the CBL1 localization sequences was expressed in E. coli BL21(DE3) and purified by NiNTA-cellulose chromatography: 1, purified sensor protein; 2, molecular mass standards. **(C)** Spectral characteristics of the modified Ca^2+^ sensor. Fluorescence emission of the purified sensor was examined in the presence (colored lines) and absence (broken black lines) of 1 mM free Ca^2+^.

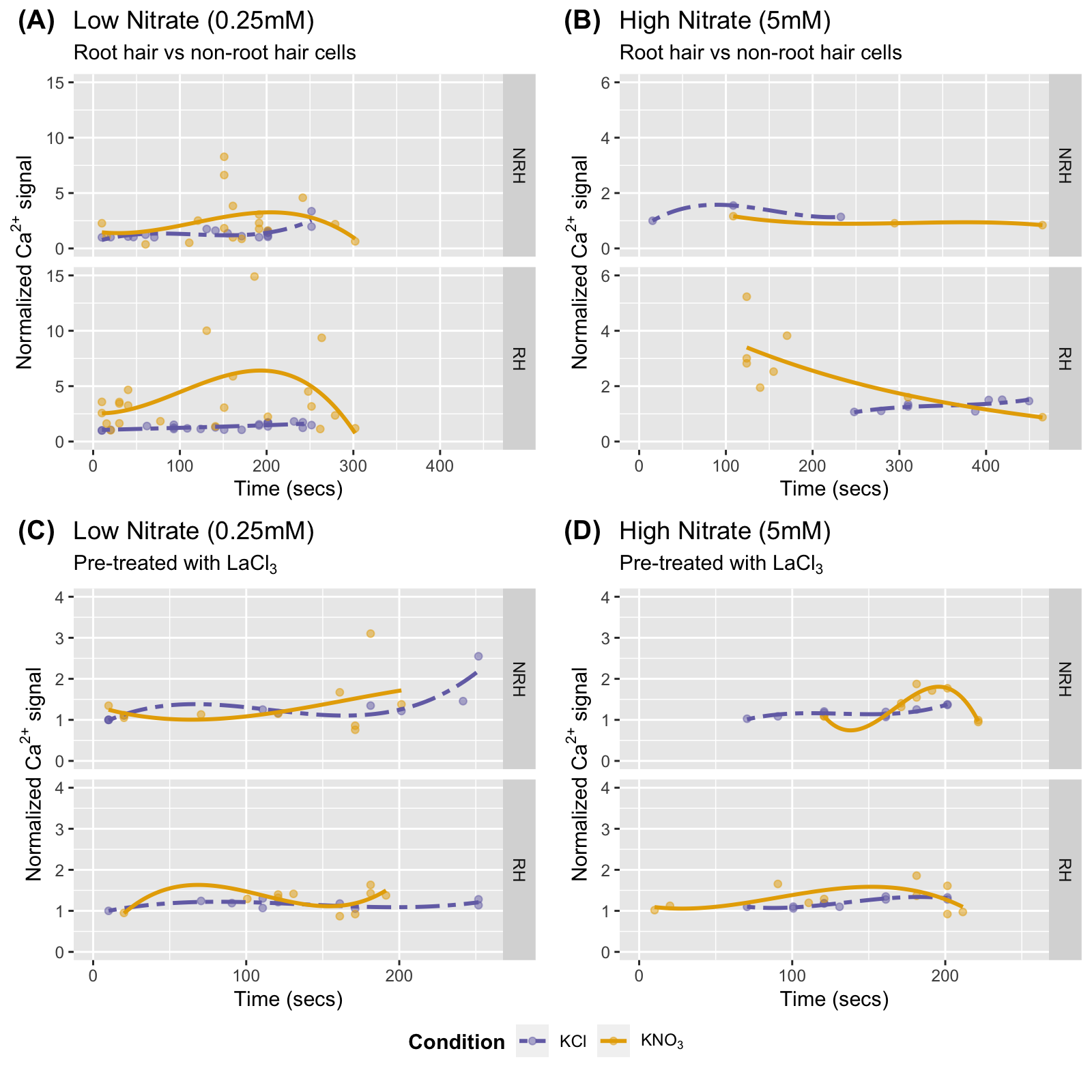
**Figure S2.** Nitrate (NO_3_^-^) kinetics showing the peak response in normalized calcium (Ca^2+^) signal at both low (0.25 mM) and high (5 mM) levels for root hairs (RH; bottom panel in each pair) compared to non-root hair epidermal cells (NRH; top panel in each pair). The median time to peak normalized Ca^2+^ signal intensity was captured around 140 sec in RHs for both low- and high- NO_3_^-^ treatments, and around 180 seconds for LaCl3 pretreated seedlings in response to low- and high- NO_3_^-^ treatments. **(A)** Variation in normalized Ca^2+^ signal in response to low (0.25 mM, n= 10 seedlings) NO_3_^-^ treatment **(B)** Variation in normalized Ca^2+^ signal in response to high (5 mM, n=2 seedlings) NO_3_^-^. **(C)** Variation in normalized Ca^2+^ signal in LaCl_3_ pretreated seedlings in response to low (0.25 mM, n= 5 seedlings) NO_3_^-^ treatment **(D)** Variation in normalized Ca^2+^ signal in in LaCl_3_ pretreated seedlings in response to high (5 mM, n=5 seedlings) NO_3_^-^ treatment.

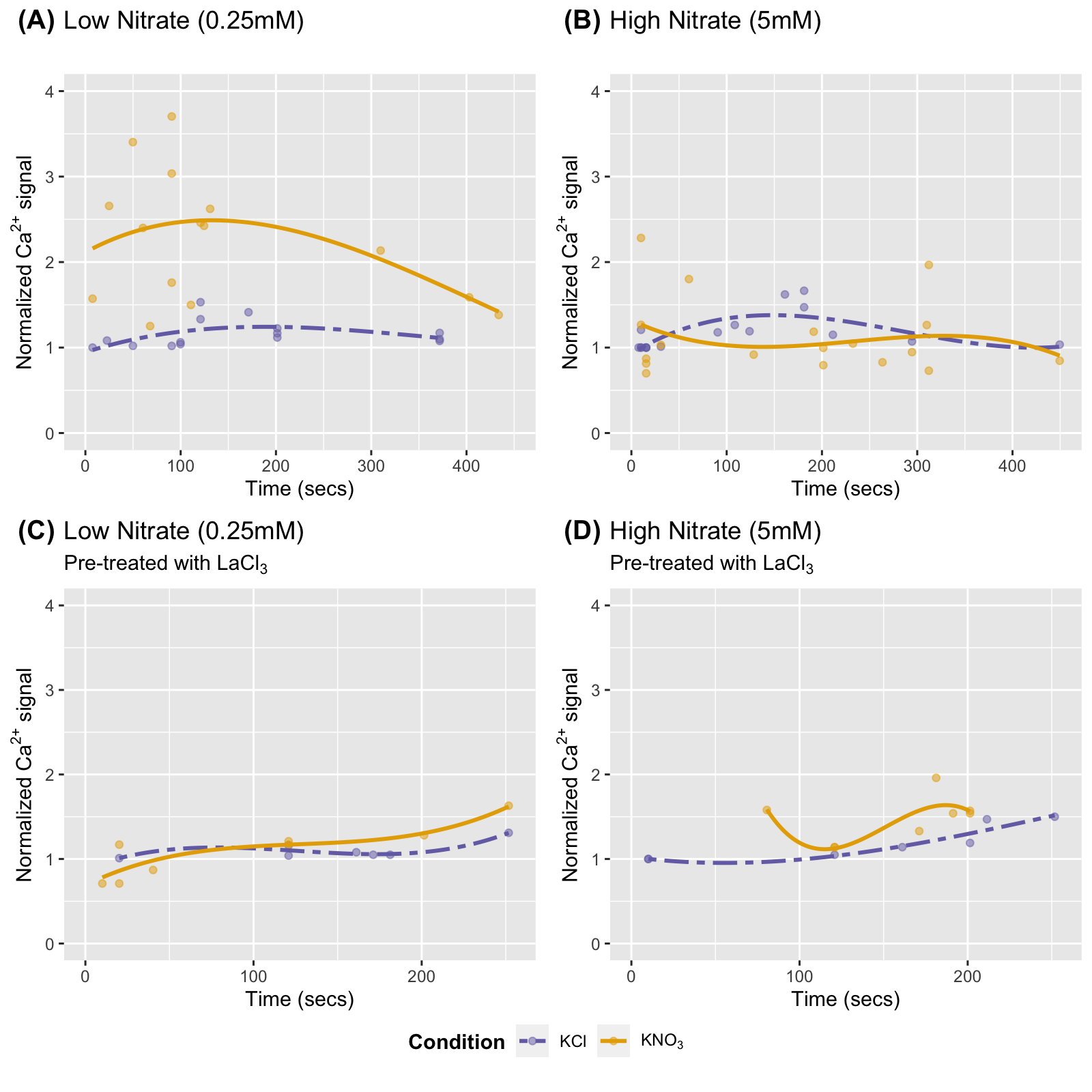

**Figure S3.** Kinetics of nitrate (NO_3_^-^) induced peak normalized calcium (Ca^2+^) signal showing an increase in low (0.25 mM) nitrate (NO_3_^-^) treatment but not in high (5 mM) treatment in bulk epidermal cells of the maturation zone. For pretreatment, the seedlings were incubated with 5mM LaCl_3_ for 1 hour prior to NO_3_^-^ treatments. **(A, B)** Variation in peak normalized Ca^2+^ signal in response to low (0.25 mM, n= 5 seedlings) NO_3_^-^ and high (5 mM, n=7 seedlings) NO_3_^-^ treatment. **(C, D)** Variation in the peak normalized Ca^2+^ signal in response to low (0.25 mM, n= 6 seedlings) NO_3_^-^ and high (5 mM, n=7 seedlings) NO_3_^-^ treatment LaCl_3_ pretreated seedlings.

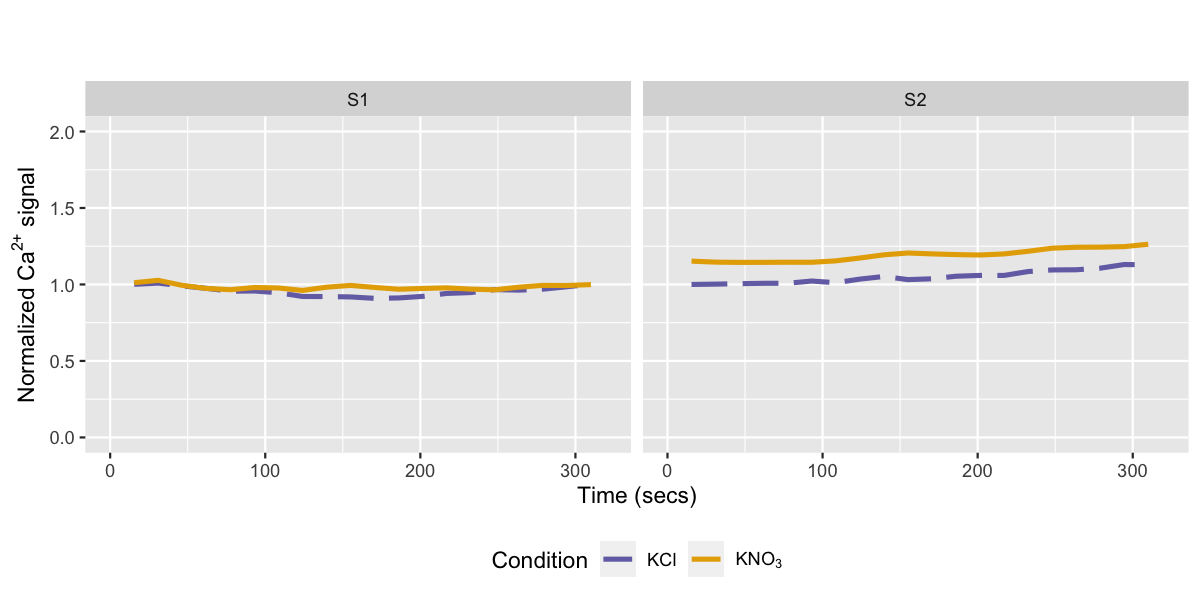

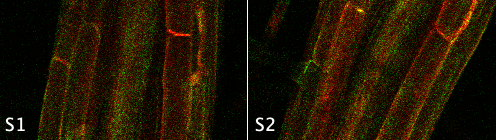

**Figure S4.** Asynchronous normalized Ca^2+^ signal in the root sections in response to the high NO_3_^-^ treatment. A representative root section for the high NO_3_^-^ treated seedling is shown with the normalized Ca^2+^ signal, where section S1 was distal to the root tip while S2 was proximal to the root tip. The root tissue imaged above shows the apparent Ca^2+^ signal through GFP fluorescence with mRuby2 RFP background and captures the response 90 seconds after high NO_3_^-^ treatment. The peak normalized Ca^2+^ signal intensity was captured around 30 seconds for section S1 and 310 seconds for S2, with 1.1- and 1.25-fold increase compared to high KCl treatment.
